# Comparing the value of mono- versus coculture for high-throughput compound screening in hematological malignancies

**DOI:** 10.1101/2022.02.18.481065

**Authors:** Sophie A. Herbst, Vladislav Kim, Tobias Roider, Eva C. Schitter, Peter-Martin Bruch, Nora Liebers, Carolin Kolb, Mareike Knoll, Junyan Lu, Peter Dreger, Carsten Müller-Tidow, Thorsten Zenz, Wolfgang Huber, Sascha Dietrich

**Affiliations:** Department of Medicine V, Hematology, Oncology and Rheumatology, Heidelberg University Hospital, Heidelberg, Germany; European Molecular Biology Laboratory (EMBL), Heidelberg, Germany; Molecular Medicine Partnership Unit (MMPU), Heidelberg, Germany; Department of Translational Medical Oncology, National Center for Tumor Diseases (NCT) Heidelberg and German Cancer Research Center (DKFZ), Heidelberg, Germany; Faculty of Biosciences, University of Heidelberg, Heidelberg, Germany; Department of Hematology and Oncology, University Hospital Düsseldorf, Düsseldorf, Germany; Department of Medical Oncology and Hematology, University Hospital Zürich, Zürich, Switzerland

## Abstract

Large-scale compound screens are a powerful model system for understanding variability of treatment response and for discovering druggable tumor vulnerabilities of hematological malignancies. However, as mostly performed in a monoculture of tumor cells, these assays disregard modulatory effects of the *in vivo* microenvironment. It is an open question whether and to what extent coculture with bone marrow stromal cells could improve the biological relevance of drug testing assays over monoculture. Here, we measured ex vivo sensitivity of 108 primary blood cancer samples to 50 drugs in monoculture and in coculture with bone marrow stromal cells. Stromal coculture conferred resistance to 52 % of compounds in chronic lymphocytic leukemia (CLL) and to 36% of compounds in acute myeloid leukemia (AML), including chemotherapeutics, BCR inhibitors, proteasome inhibitors and BET inhibitors. While most of the remaining drugs were similarly effective in mono- and coculture, only the JAK inhibitors ruxolitinib and tofacitinib exhibited increased efficacy in AML and CLL stromal coculture. We further confirmed the importance of JAK-STAT signaling for stroma-mediated resistance by showing that stromal cells induce phosphorylation of STAT3 in CLL cells. We genetically characterized the 108 cancer samples and found that drug-gene associations agreed well between mono- and coculture. Overall, effect sizes were lower in coculture, thus more drug-gene associations were detected in monoculture than in coculture. Our results suggest a two-step strategy for drug perturbation testing, with large-scale screening performed in monoculture, followed by focused evaluation of potential stroma-mediated resistances in coculture.

## Introduction

*Ex vivo* compound screening has improved our understanding of the phenotypic and molecular heterogeneity of tumor diseases [1–10]. In patients with hematological malignancies, profiling drug responses on demand has even been demonstrated to support clinical decision making by suggesting personalized treatment options [11, 12]. Most of these studies face the problem that, deprived of microenvironmental stimuli, leukemia cells undergo spontaneous apoptosis *ex vivo* [13, 14]. There are several approaches for modeling the leukemia microenvironment *ex vivo,* for instance by adding conditioned medium from stromal cells [15, 16] or by providing specific stroma-secreted cytokines [17]. However, not only soluble factors, but also the direct contact with stromal cells play an essential role in promoting the survival of leukemia cells in the bone marrow [18]. Coculture studies revealed that bone marrow-derived stromal cells protect leukemia cells even from drug-induced apoptosis [19–22], which may contribute to residual disease [23] and the emergence of resistant clones [24]. Therefore, stroma-leukemia coculture models are considered a potential *ex vivo* platform to profile drug responses of tumor cells while mimicking the interactive effects of the microenvironment [10, 20, 25–27].

Though coculture models appear more natural to profile drug response *ex vivo*, given the complexity and extra effort to establish and read-out such a model, the application must be carefully considered. Unfortunately, the validity of coculture models has not been tested rigorously, and current evidence is limited to individual compounds probed in small-scale coculture studies [19–21, 28–35].

To systematically assess whether coculture studies provide superior biological insights, we performed a large-scale study comparing compound efficacy in leukemia monoculture and leukemia-stroma coculture. We used the well-established bone-marrow derived stroma cell line HS-5 and an imaging-based platform to investigate not only drug effects in mono- and leukemia-stroma coculture but also to capture cellular changes due to the stromal environment and drug treatments. Finally, we suggest a two-stage strategy of high-throughput drug perturbation in monoculture followed by targeted evaluation of stroma-mediated resistance in cocultures.

## Materials and methods

### Cell culture

HS-5 cells were cultured in DMEM (Thermo Fisher Scientific) supplemented with 10 % fetal bovine serum (FBS; Thermo Fisher Scientific), 1 % penicillin/streptomycin (Thermo Fisher Scientific) and 1 % glutamine (Thermo Fisher Scientific) in a humidified atmosphere at 37° C and 10 % CO2.

### Patient samples

Written consent was obtained from all patients according to the declaration of Helsinki. Also, our study was approved by the Ethics Committee of the University of Heidelberg. Samples were selected based on availability and tumor cell content higher than 80 %. Clinical flow cytometry data were used to estimate the proportion of malignant cells in collected blood samples. Peripheral blood mononuclear cells (PBMCs) were isolated using Ficoll density gradient centrifugation. Cells were viably frozen in RPMI (Thermo Fisher Scientific) containing 45 % FBS (Thermo Fisher Scientific) and 10% DMSO (SERVA Electrophoresis GmbH) and kept on liquid nitrogen until use. Cells were thawed freshly before the experiment and rolled in serum containing medium for 3 hours on a roll mixer at room temperature to allow cells to recover. To deplete dead cells, which form clumps during this procedure, the suspension was filtered through a 40 μm cell strainer (Sarstedt). Cell viability and counts were analyzed using Trypan Blue (Thermo Fisher Scientific). Percentages of alive cells always exceeded 90 % at culture start or freezing of pellets.

### IGHV status analysis

For the analysis of IGHV status RNA was isolated from 1×10^7^ PBMCs and cDNA was synthesized via reverse transcription. Subsequent PCR reactions and analyses were performed according to Szankasi and Bahler with minor modifications [36]. A detailed description can be found in the supplementary methods of this manuscript.

### Panel sequencing of CLL samples

We performed an analysis of gene mutations of the CLL candidate genes NOTCH1, SF3B1, ATM, TP53, RPS15, BIRC3, MYD88, FBXW7, POT1, XPO1, NFKBIE, EGR2 and BRAF. A detailed description of the analysis can be found in the supplementary methods of this manuscript.

### DNA copy number variants

Assessment of DNA copy numbers was done using Illumina CytoSNP-12 and HumanOmni2.5-8 microarrays and read out using an iScan array scanner. Fluorescence in situ hybridization (FISH) analysis was performed for del11q22.3, del17p13, del13q14, trisomy 12, gain8q24 and gain14q32. Only alterations present in at least three patients and absent in at least three patients were considered.

### Drug plate preparation

For the screen, 50 drugs were probed at 3 different concentrations (Supplementary Table 1). Drug concentrations ranged from subnanomolar to low micromolar and were chosen based on previous experience with the drugs [3]. Increase of the concentration was 15-fold per step to cover a broad spectrum of concentrations. Drugs were diluted according to the manufacturer’s instructions. Further dilution was carried out in DMSO (SERVA Electrophoresis GmbH) and master plates containing 4 μL of diluted drugs were frozen at −20 °C for direct use on the screening days.

### Compound screening of mono- and cocultures

Drug screens were carried out in CellCarrier-384 Ultra Microplates (Perkin Elmer) with a seeding density of HS-5 stromal cells of 1×10^4^ cells/well and 2×10^4^ patient cells per well. The screen was carried out in DMEM (Thermo Fisher Scientific) supplemented with 10 % human serum (male AB, H6914-100ml Batch SLBT2873, Sigma-Aldrich), 1 % penicillin/streptomycin (Thermo Fisher Scientific) and 1 % glutamine (Thermo Fisher Scientific) at a final volume of 40 μL in the culture plates. Cells were incubated at 37 °C in a humidified atmosphere and 10 % CO2 for 3 days. A detailed description of the screen can be found in the supplementary methods section.

### Spinning disk confocal microscopy

High-throughput screening was conducted using Opera Phenix High Content Screening System (Perkin Elmer). CLL screening plates were stained with 4 μg/ml Hoechst 33342 (Invitrogen) and 1 μl/ml lysosomal dye NIR (Abcam). Plates of non-CLL entities were additionally stained with Calcein AM (1 μM, Invitrogen). Three positions per well with a stack of ten images at a distance of 1.2 μm were acquired with a 40x water objective in confocal mode.

### Primary mesenchymal stromal cells cocultures

Drug screen results for 1.5 μM JQ1, 0.6 μM Fludarabine, 22.5 μM tofacitinib and 9 μM ruxolitinib were validated in cocultures with primary mesenchymal stromal cells (MSCs) derived from three different healthy donors. Each condition was assessed in technical duplicates. For a detailed description see the supplementary methods of this manuscript.

### Processing of images (CLL)

Images of CLL samples were processed using the image analysis software Harmony (Perkin Elmer). Results were further analyzed in the statistical programming language R (R Core Team, 2018). For a detailed description see the supplementary methods of this manuscript.

### Image analysis in non-CLL entities

Maximum intensity projection and gamma correction (gamma = 0.3) was applied to all images. All 3 color channels (lysosomal dye, Calcein and Hoechst) were combined to generate RGB overlays. Each image (2160 x 2160, omitting the color channel axis) was cut into 9 blocks of size 720 x 720 to speed up training and prediction.

Faster R-CNN object detection model [37] with Inception v2 [38] backbone architecture was used to detect patient-derived leukemia and lymphoma cells. The two defined classes were viable and apoptotic leukemia cells. The object detection model implemented in TensorFlow 1.14 was trained for 21,000 epochs on coculture images from 5 AML samples. 5 control and 5 drug-treated well images were randomly selected from each of the five AML plates, resulting in 5 * 10 * 9 = 450 images that were split into train / test sets with 80% / 20% ratio. The average precision (AP) on the test set was 0.99 and 0.93 for viable and apoptotic leukemia cells, respectively. The area under the ROC curve (AUCROC) was 0.98 for both viable and apoptotic leukemia cells.

### Morphological profiling, quality control and normalization

After image segmentation, morphological properties describing size, intensity, shape, and texture were computed for each cell. Morphological profiling of patient-derived leukemia cells produced 1401 image features in non-CLL entities and 934 features in CLL. In all downstream analyses, we used only a subset of features with high replicate correlation (r > 0.5). After filtering based on replicate correlation, we obtained 173 morphological features in non-CLL entities and 194 features in CLL. All morphological properties were normalized to control values. Mean and standard deviation of each image feature were estimated using untreated wells in mono- and coculture, respectively. All morphological features were centered and scaled:

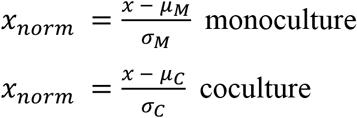

### Spontaneous apoptosis, drug sensitivity and normalization

Only viable and apoptotic leukemia cell counts were used for drug sensitivity analysis. Viability was computed as the ratio of viable cell count to the total cell count. For each sample, baseline viabilities (*b_M_,b_C_*) were defined as mean viabilities of untreated wells of the respective culture condition. Untreated wells on the plate edge were excluded, resulting in 11 and 13 wells used for estimation of baseline viability in mono- and coculture, respectively. Spontaneous apoptosis rate was evaluated as the complement of baseline viability:

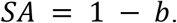

Drug sensitivities were computed by normalizing viabilities to baseline values of the respective culture condition:

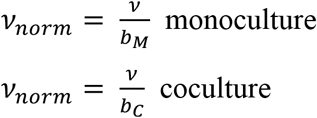

### Compound efficacy changes in coculture

For each drug, we selected the concentration with maximum variance in terms of normalized viability and applied a paired *t*-test with the null hypothesis H_0_ assuming equal drug sensitivities in mono- and coculture. Drug concentrations toxic to stromal cells were excluded prior to statistical testing but were retained for dose-response fitting.

To compute the effect size, median dose-response curves were computed for mono- and coculture. The effect size was calculated as the percentage change in area under the dose-response curves in coculture:

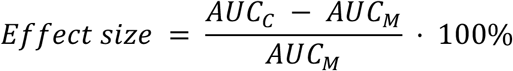

In CLL coculture, compounds with changed efficacy had adjusted p-values < 0.01 and |effect size| > 5 %. In AML coculture, the same thresholds were used, except for those compounds that change efficacy in CLL coculture, for which only the effect size cutoff of 5% was used.

### Drug-gene associations

For 80 CLL samples, genetic features such as IGHV mutation status, somatic mutations (TP53, ATM, etc.) and chromosomal aberrations (del11q, trisomy 12, etc.) were available. To test whether mean drug sensitivities of wildtype and mutated cases were equal, we applied a *t*-test on normalized viabilities for each drug stratified by mutational status. The statistical tests were performed separately in mono- and coculture.

### Western blot analysis

To assess the impact of stroma coculture on STAT3 phosphorylation in CLL cells, DMEM medium supplemented with 10% human serum (male AB, H6914-100ml Batch SLBT2873, Sigma-Aldrich), 1 % penicillin/streptomycin (Thermo Fisher Scientific) and 1 % glutamine (Thermo Fisher Scientific) or 5×10^6^ HS-5 cells were pre-plated into 10 cm dishes. After 3 hours CLL cells were added at 1.5×10^7^ cells/dish to establish mono- and cocultures. DMSO (0.22 %; SERVA Electrophoresis GmbH), ruxolitinib (10 μM) or tofacitinib (22 μM) were added. After incubation for 48 hours CLL cells were carefully harvested. Cells were counted using Trypan Blue (Thermo Fisher Scientific) and contamination with HS-5 cells was excluded by visual inspection. To assess the impact of soluble factors produced by stroma, HS-5 cells or primary MSCs were cultured in DMEM medium supplemented with 10% FBS (Thermo Fisher Scientific), 1 % penicillin/streptomycin (Thermo Fisher Scientific) and 1 % glutamine (Thermo Fisher Scientific) or Bulletkit medium (Lonza) respectively. Conditioned medium was harvested after 3 days of culture. After the removal of cellular debris by centrifugation at 1000 g, aliquots of conditioned medium were frozen. 7.5×10^6^ CLL patient cells in DMEM medium supplemented with 10 % FBS (Thermo Fisher Scientific), 1 % penicillin/streptomycin (Thermo Fisher Scientific), 1 % glutamine (Thermo Fisher Scientific) and 25 % conditioned medium were seeded into 10 cm dishes. Cells were harvested after culturing for 48 hours. Western Blot was performed using the primary antibodies anti-phospho-STAT3^Tyr705^ (Cell Signaling Technology, #9145), anti-STAT3 (Cell Signaling Technology, #30835), anti-β-actin (Proteintech Group, #66009-1-Ig), and the secondary antibodies anti-mouse-IgG-HRP-conjugated (Proteintech Group, #SA00001-1), and anti-rabbit-IgG-HRP-conjugated (Proteintech Group, #SA00001-2). A detailed description of the protocol can be found in the supplementary methods of this manuscript.

## Data availability

Primary imaging data will be available via Imaging Data Resource (IDR) upon publication. Raw and normalized drug response data of mono- and coculture are available on GitHub (https://github.com/vladchimescu/coculture.git).

## Software availability

Image analysis and morphological profiling were conducted in Python and the code is available on Github (https://github.com/vladchimescu/microscopy-notebooks.git). Statistical analysis of processed viability and morphological feature data was performed in R and the code is available on Github (https://github.com/vladchimescu/coculture.git).

## Results

### Imaging-based compound screen in leukemia-stroma coculture

We established a microscopy-based platform (Figure 1) for compound screening in primary blood cancer cells cocultured with the HS-5 bone marrow stromal cell line [39], which has been demonstrated to reproduce most features of bone marrow-derived stromal cells [40]. Using this platform, we screened 50 compounds at three concentrations (Supplementary Table 1) in 108 leukemia and lymphoma samples (Supplementary Tables 2-3) in mono- and coculture (Figure 1), including chronic lymphocytic leukemia (CLL, n = 81), acute myeloid leukemia (AML, n = 17), T-cell-prolymphocytic leukemia (T-PLL, n = 4), mantle cell lymphoma (MCL, n = 4) and hairy cell leukemia (HCL, n = 2). An exposure time of 72 hours and drug concentrations aiming for high, medium, or low toxicity were selected based on a previous internal high-throughput compound screen [3]. After 72 hours, we used Hoechst to stain nuclei in all samples and employed confocal microscopy to readout viability and morphological changes in leukemia and lymphoma cells. As previously described, viable CLL cells were identified based on the Hoechst-stained nucleus area [41] (Supplementary Figure 1A), whereas in non-CLL entities an additional staining of the cytoplasm using Calcein was required to distinguish viable and dead cells (Supplementary Figure 1B, see Method section for details). Our primary readout was viability, defined as the viable fraction of leukemia cells. To adjust for spontaneous apoptosis, viabilities in drug-treated wells were normalized to viabilities in untreated wells. The viability readout of our platform was highly reproducible between replicates with correlations of *R* = 0.88 in coculture and R = 0.92 in monoculture (Supplementary Figure 1C), and between Hoechst- and Calcein-based readout in CLL samples with correlations of R = 0.92 (Supplementary Figure 1D).

**Figure 1.**
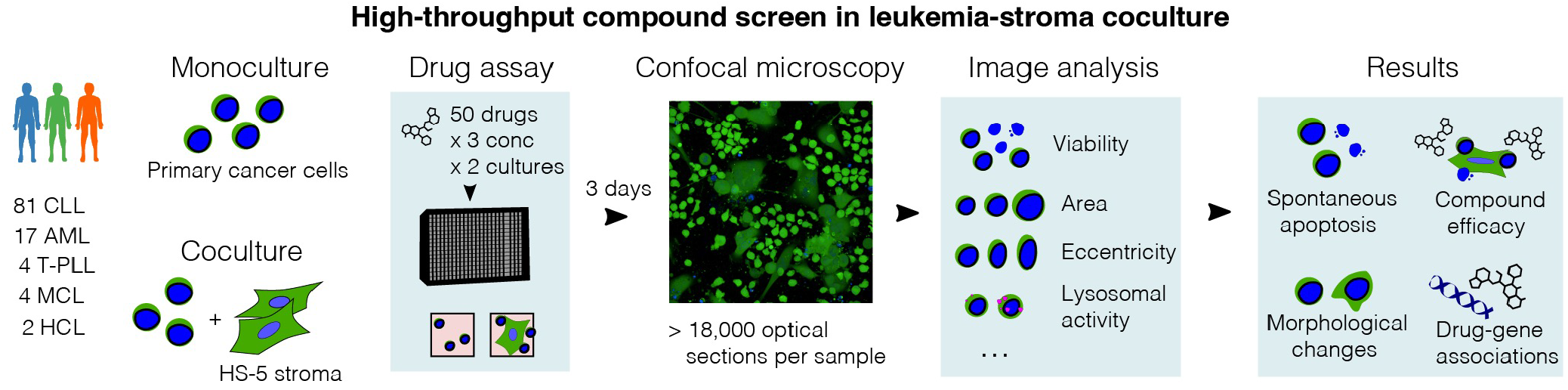
Imaging-based coculture screen in primary leukemias and lymphomas. Study outline. 50 compounds were probed in 108 primary leukemia and lymphoma samples. Confocal microscopy images of leukemia cells alone (in monoculture) and in coculture with the HS-5 stromal cell line were acquired to compute viability and morphological properties.

### Degree of stromal protection varies across probed compounds

To assess the degree of spontaneous apoptosis, we determined median raw viability of untreated wells. In monoculture, proportions of viable leukemia cells in the absence of drug treatment were highly variable, ranging from 10% to over 90% (Supplementary Figure 2). Interestingly, samples with low monoculture viability (< 50% alive cells) showed the highest increase in viability in coculture (Supplementary Figure 2), reflecting their stronger dependence on the microenvironment signals.

Next, we determined leukemia and lymphoma cell viability after *ex vivo* exposure to fifty different compounds and compared normalized viabilities in monoculture (Supplementary Figure 3) with those in coculture (Supplementary Figure 4) for each compound using a paired t-test. 26 out of 50 (= 52 %) compounds in CLL-stroma coculture and 18 out of 50 (= 36 %) compounds in AML-stroma coculture were less toxic compared to their corresponding monoculture conditions (Figure 2A, Supplementary Table 4). Quantitative assessment of drug efficacy changes in coculture revealed similar patterns in AML and CLL (Figure 2A, Supplementary Figure 5). In line with previously reported findings [19, 20, 28, 42–44], coculture significantly reduced the toxicity of the chemotherapeutics (fludarabine, doxorubicin, cytarabine) both in CLL and AML (Figure 2A). Likewise, the proteasome inhibitors carfilzomib and ixazomib, as well as the BET inhibitors JQ1 and I-BET-762, showed significantly reduced efficacy in CLL and AML cocultures compared to monocultures (Figure 2A). By coculturing CLL cells with primary mesenchymal stromal cells (MSCs), we reproduced stroma-mediated protection against drug-induced apoptosis using fludarabine as an example (Figure 2B). Unlike HS-5 cells, primary MSCs have not undergone immortalization and were subjected only to a limited time of *ex vivo* culturing. Similarly, we used primary MSCs to confirm the protection against BET inhibitor-mediated toxicity in CLL (Figure 2C). In contrast, we identified a considerable proportion of drugs that were similarly effective in CLL (44 %) and AML (58 %) cocultures (Figure 2A, Supplementary Table 4). Among these were both clinically relevant drugs, such as BCR-Abl/Src inhibitor dasatinib, FLT3 inhibitor quizartinib, CDK inhibitor palbociclib (Figure 2A), and several experimental compounds such as Mdm2 inhibitor nutlin 3a, BH3 mimetics obatoclax mesylate and UMI-77, Akt inhibitor MK2206, and NFkB inhibitors EVP4593 and BAY11-7085 (Figure 2A). These results suggest that the bone marrow microenvironment selectively influences the efficacy of many but not all compounds.

**Figure 2.**
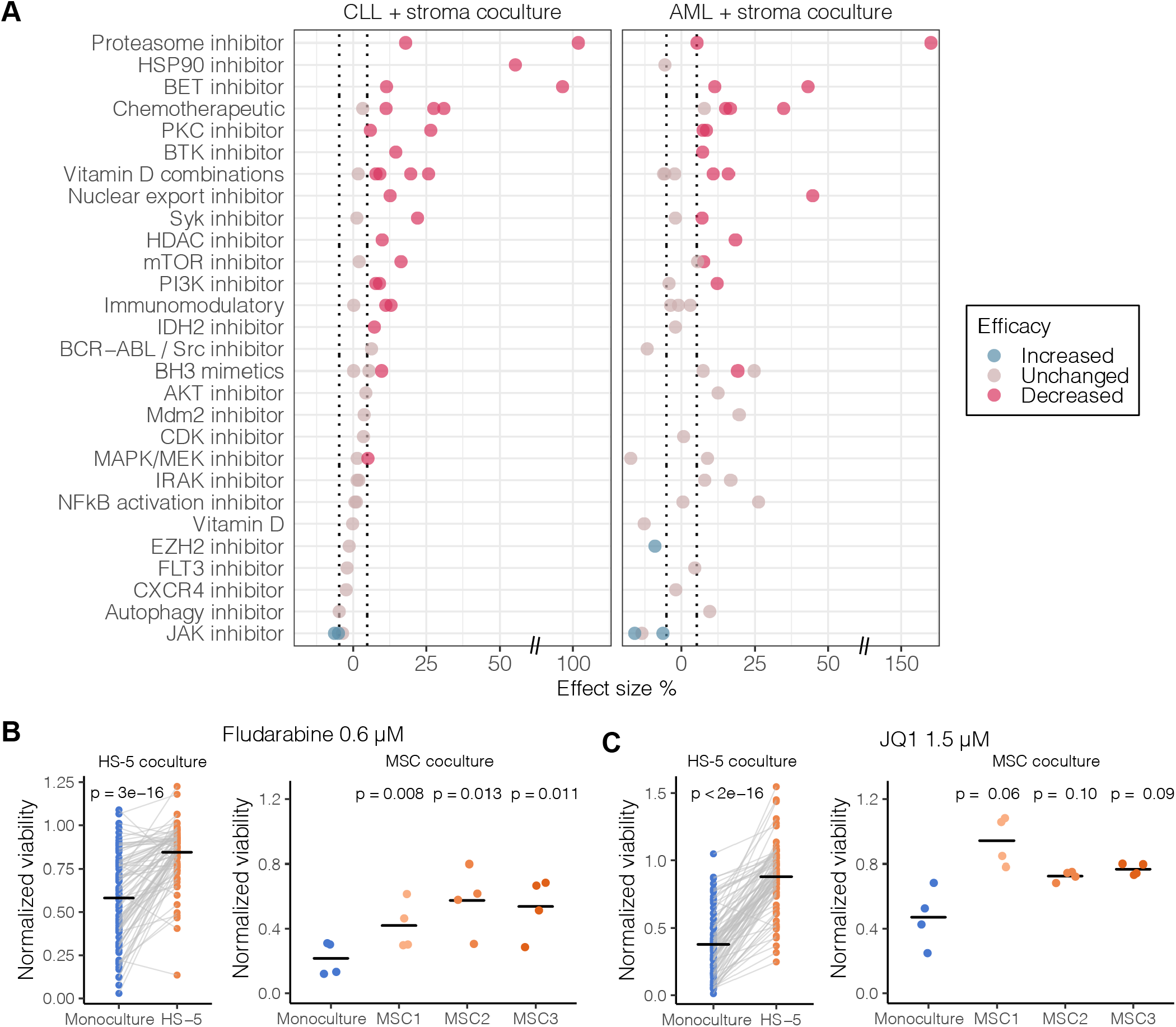
Stroma-mediated modulation of compound efficacy. A) Compounds ordered by drug class and colored by their change in efficacy in CLL-stroma and AML-stroma coculture compared to monoculture. The x-axis shows the effect size of the drug response change relative to monoculture (see Materials and Methods). T-test was used to assess changes of compound efficacy in coculture (FDR = 0.01). The effect size threshold of 5% is indicated as dotted vertical lines. B-C) Validating the effect of fludarabine (B) and JQ-1 (C) from the HS-5 coculture screen in cocultures of CLL with primary mesenchymal stromal cells (MSCs). P values compare the coculture mean with the reference value in monoculture. MSC1, MSC2, MSC3 were derived from 3 healthy donors.

### Stroma-leukemia coculture increases toxicity mediated by JAK inhibitors

Among all compounds, only the JAK inhibitors tofacitinib and ruxolitinib were more effective in CLL and AML coculture than in monoculture (Figure 2A). Again, we confirmed this effect by coculturing CLL cells with primary MSCs and exposing them to ruxolitinib (Figure 3A) and tofacitinib (Figure 3B). Importantly, the JAK-STAT pathway has been suggested as a key mediator of stromal protection [31, 32, 45, 46]. Indeed, we observed that the presence of bone marrow stromal cells increased phosphorylation of STAT3 at Tyr705 in CLL cells, which could be reversed by simultaneous exposure to JAK inhibitors (Figure 3C). Conditioned medium from HS-5 cells or primary MSCs was sufficient to increase STAT3 phosphorylation (Figure 3D), demonstrating that JAK-STAT-mediated protection is based on the exchange of soluble factors. These results highlight the importance of targeting components of the soluble microenvironment for disrupting the interaction between stromal and leukemia cells.

**Figure 3.**
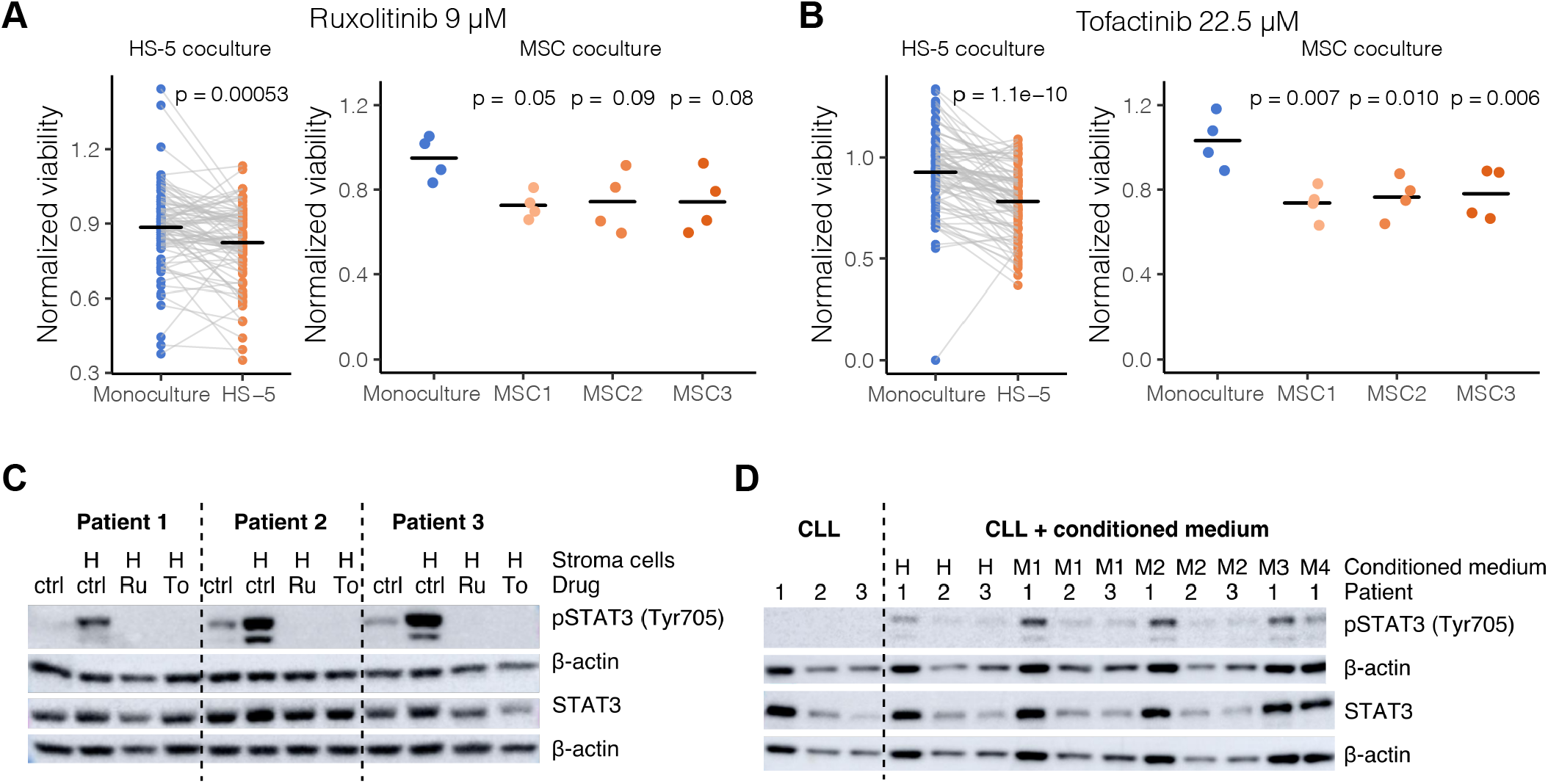
Stroma-leukemia coculture increases toxicity mediated by JAK inhibitors. A-B) Validating effect of ruxolitinib (A) tofacitinib (B) from the HS-5 coculture screen in cocultures of CLL with primary mesenchymal stromal cells (MSCs). P values compare the coculture mean with the reference value in monoculture. MSC1, MSC2, MSC3 were derived from 3 healthy donors. C) STAT3 was phosphorylated in CLL cells cocultured with HS-5 cells. STAT3 phosphorylation could be reversed by inhibition with ruxolitinib or tofacitinib. D) STAT3 was phosphorylated in CLL cells in the presence of conditioned medium derived from stromal cells. Ctrl = solvent control (DMSO), Ru = ruxolitinib (10 μM), To = tofacitinib (22 μM). H = cocultures with HS-5 cells, M1-4 = cocultures with MSC cells from four different donors.

### Coculture recapitulates most clinically established drug-gene associations

To identify and compare drug-gene associations between mono- and coculture, we characterized key genetic features of CLL samples, including TP53 mutation, IGHV status, and trisomy12 status. For each drug-gene pair we performed a *t*-test, comparing drug responses in wildtype and mutated groups, as shown for nutlin 3a and TP53 mutation or ibrutinib and IGHV status (Figure 4A). The comparison of the *t*-statistic values in mono- and coculture are summarized in Figure 4B, with significant associations (FDR < 0.1) highlighted. While the direction of drug-gene associations was preserved in CLL coculture (Figure 4B, Supplementary Figure 6), we observed that associations of BCR inhibitors with IGHV and trisomy12 status exhibited smaller effect sizes in CLL coculture than in monoculture (Figure 4C). Consequently, some well-established associations, such as the increased sensitivity of the U-CLL group to ibrutinib [3, 47], could be detected in monoculture but did not reach statistical significance in coculture (Figure 4A, Supplementary Table 5). In line with that, we observed that stroma-mediated protection from BCR inhibitors was stronger in U-CLL than in M-CLL samples (Figure 4D). Also, trisomy12 positive samples treated with BCR inhibitors were better protected by the stromal microenvironment than trisomy12 negative samples (Figure 4D).

**Figure 4.**
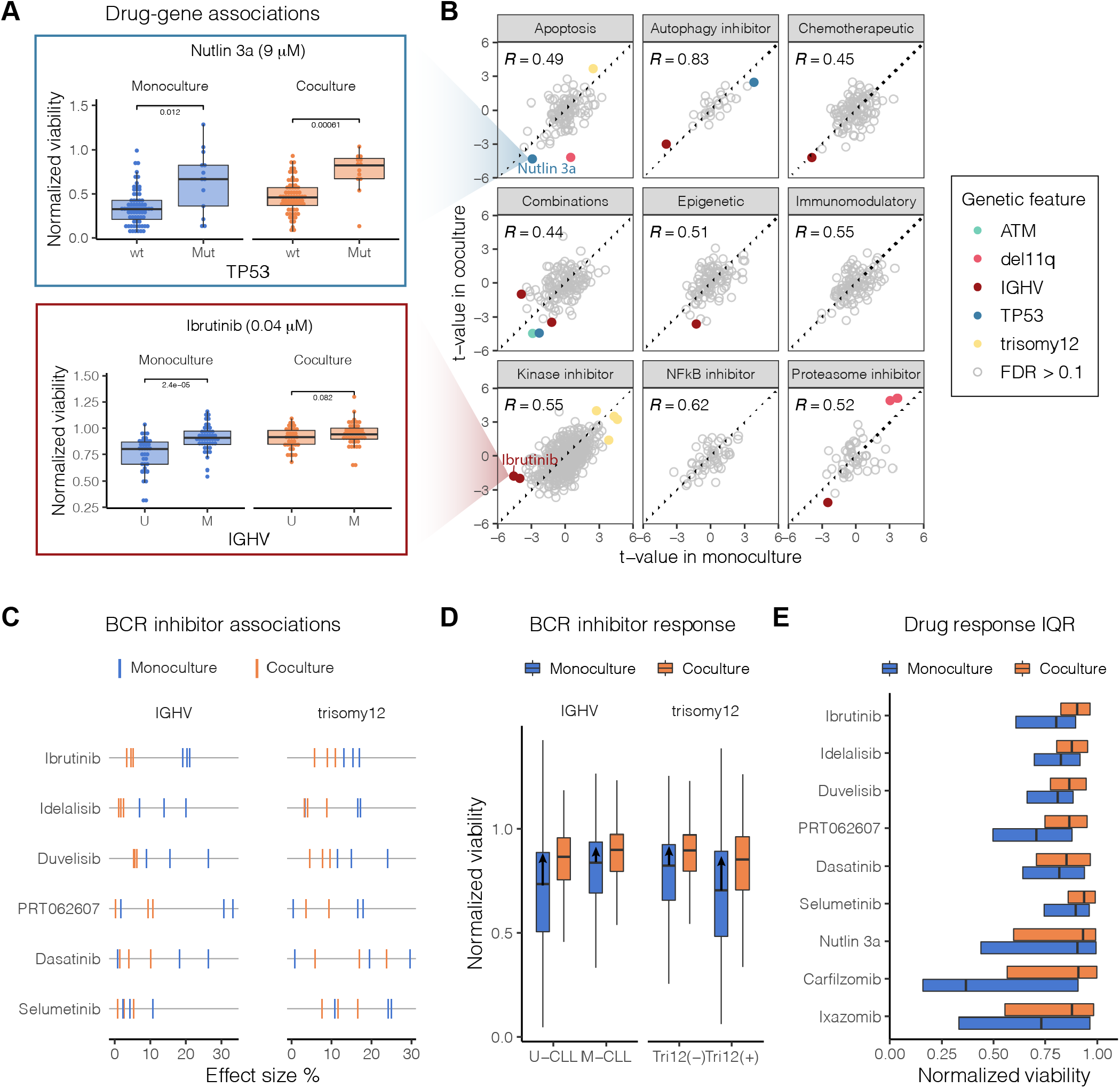
Drug-gene associations in coculture. A) Boxplots showing response to nutlin 3a and ibrutinib stratified by culture condition and mutational status. B) Comparison of druggene association statistics in mono- and coculture. The x- and y-axes show the t-statistic values of drug-gene associations in mono- and coculture at a given concentration. C) Effect size of IGHV and trisomy 12 associations with B cell receptor (BCR) inhibitor response. The tick marks, colored by culture condition, show the absolute value of the effect size at 3 probed drug concentrations. D) The boxplots, colored by culture condition, show BCR inhibitor response stratified by IGHV mutational status (U-CLL / M-CLL) and trisomy12 (negative / positive). The arrows indicate differences between mono- and coculture medians, i.e., viability gain in coculture. E) Drug response variability in CLL samples treated with BCR inhibitors, nutlin 3a and proteasome inhibitors stratified by culture condition. The boxplots compare the interquartile ranges (IQR) of drug sensitivities in mono- and coculture.

Coculture reduced not only effect size estimates but also the drug response variability of many compounds, as observed for BCR inhibitors, nutlin 3a and proteasome inhibitors (Figure 4E). This variance reduction in coculture offset the decrease in effect size and thus enhanced some drug-gene associations, such as higher sensitivity of del11q positive samples to proteasome inhibitors (Figure 4B, Supplementary Table 5). Despite reduced technical variation, the number of discovered drug-gene associations was higher in monoculture. Thus, monoculture *ex vivo* drug perturbation studies represent a sensitive first-line screening approach to detect drug-gene associations.

### Image-based phenotyping provides additional insights into drug effects

Beside Hoechst, we stained all samples with a lysosomal dye aiming to measure morphological features describing nucleus and cell shape in the context of mono- and coculture. Therefore, we used reproducible morphological properties with replicate correlations *R* > 0.5 (Supplementary Figure 7, see Materials and Methods). First, we investigated the impact of the bone marrow microenvironment on unperturbed leukemia and lymphoma cells. In AML, a joint t-SNE of viable leukemia cells based on their morphological properties revealed the separation of mono- and coculture leukemia cell populations (Figure 5A). We found increased Calcein eccentricity and convex area of AML and T-PLL cells in coculture (Figure 5B, Supplementary Figure 8A), suggesting that cells of these disease entities generally take on more elongated shapes in the presence of stromal cells. For B-cell lymphoma and CLL, we did not detect any clear changes in morphology (Figure 5B, Supplementary Figure 8B).

**Figure 5.**
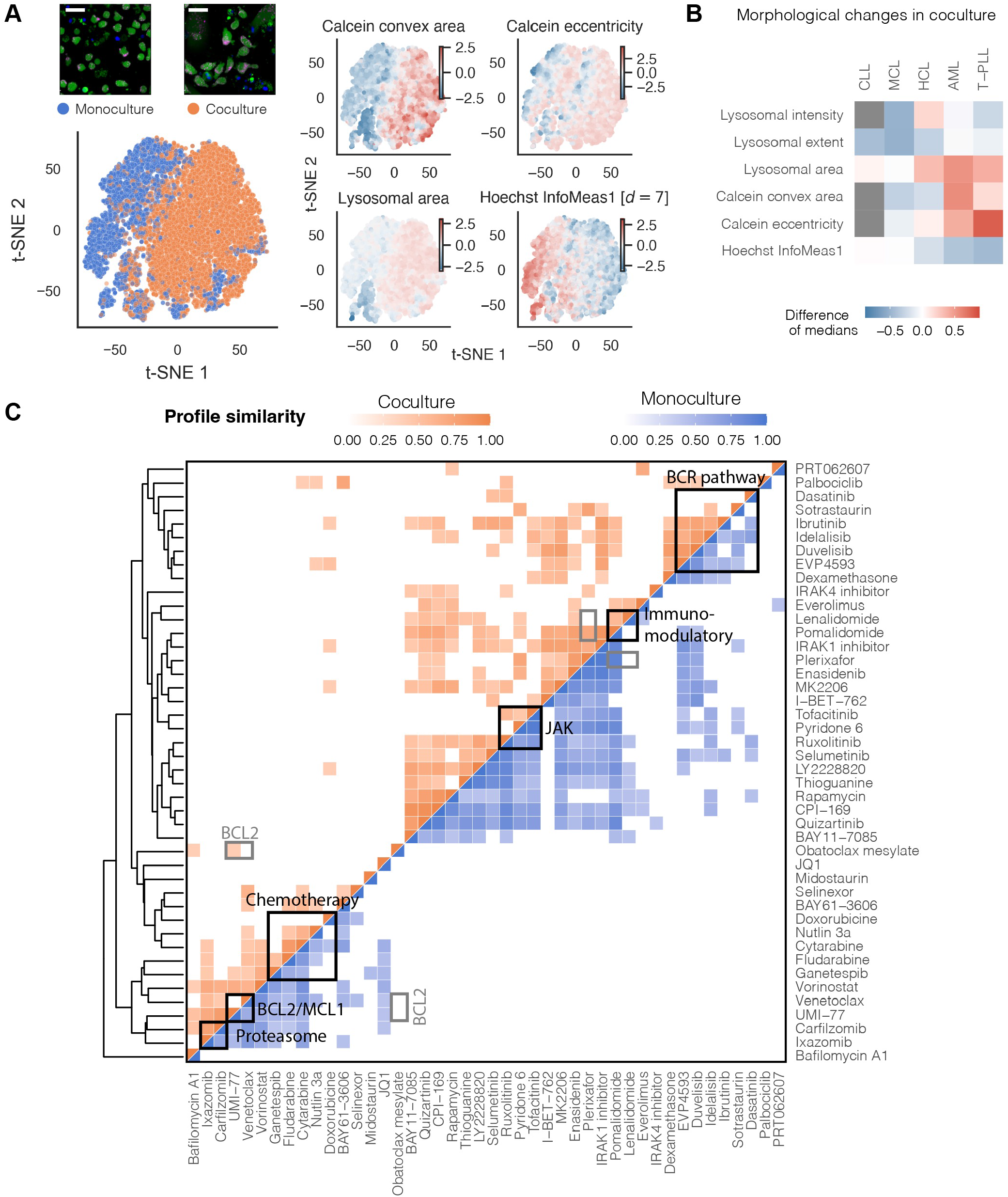
Compound similarity in mono- and coculture. A) Joint t-SNE embedding of viable leukemia cells in mono- and coculture controls of an AML sample. Coloring by morphological features revealed that AML cells in coculture had more elongated shapes (higher eccentricity), larger cell (Calcein) and lysosomal area as well as lower local correlation between pixel intensity values in x- and y-direction (Hoechst InfoMeas1). Scale bar in the left top corner is equal to 50 μm. B) Heatmap showing morphological changes in coculture controls across all screened disease entities. Grey indicates missing values. C) Aggregated compound profiles were used to generate a hierarchical clustering of all probed compounds, excluding combinations. Drug-drug correlations were computed separately in mono- and coculture. Only high correlations (r > 0.4) are indicated in the heatmap. See Method section for details.

Finally, we aggregated viability and morphological features to generate high-dimensional compound profiles of all screened compounds in mono- and coculture (Figure 5C). Hierarchical clustering recapitulated functional drug classes including BCR inhibitors, immunomodulatory imide drugs, JAK inhibitors, chemotherapeutics, BH3 mimetics, and proteasome inhibitors (Figure 5C). We observed that several drugs displayed higher similarity in monoculture. For instance, while most BCR inhibitors were strongly correlated with one another in both mono- and coculture, high correlations of sotrastaurin and dasatinib with the other BCR inhibitors were lost in coculture (Figure 5C). JAK inhibitors clustered together, with a high correlation between ruxolitinib and pyridone-6 observed only in monoculture. Likewise, the profiles of BH3 mimetics, venetoclax and UMI-77, were more similar in monoculture. Since many compounds display higher within-class heterogeneity in coculture, we hypothesize that stromal effects vary among the drugs assigned to the same functional class.

To determine relative importance of microscopy for compound profiling, we generated and compared clustering results based on image features alone and based on viabilities (Supplementary Figure 9). This revealed that the BCR inhibitor class could be recapitulated without image features, while the clustering of proteasome inhibitors or BH3 mimetics was mainly driven by morphological features (Supplementary Figure 9). This suggests that morphological profiling is useful to infer drug mode of action of certain compound classes.

### Comparison of mono- and coculture for microscopy-based screening

Our study provides a comprehensive assessment of the stroma coculture model for compound screening in hematological malignancies. To provide assistance for future compounds screening efforts, we summarized the advantages and shortcomings of coculture regarding image-based screening in Table 1.

**Table 1.**
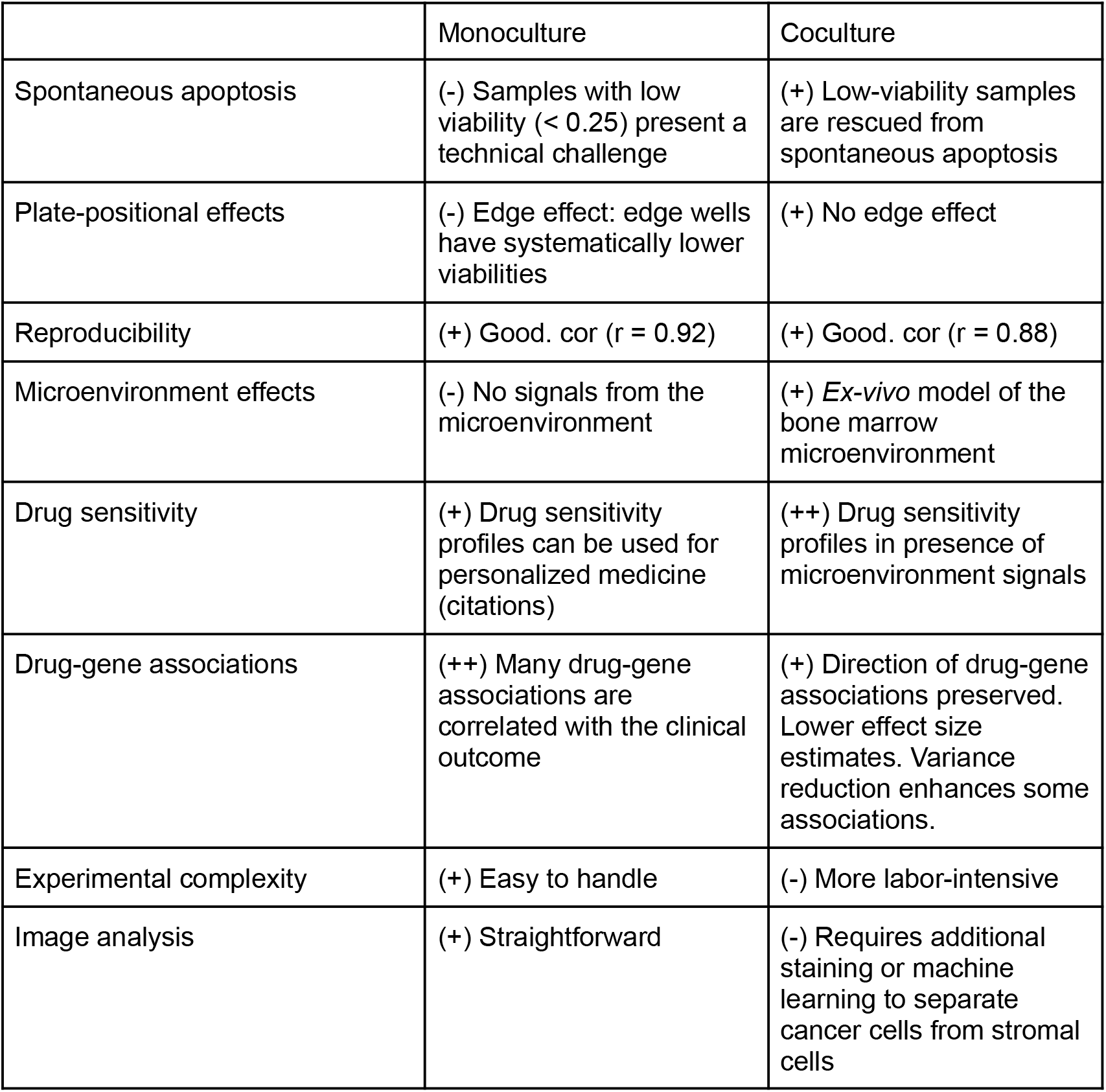
Comparison of mono- and coculture: advantages and challenges.

## Discussion

In this study, we established a microscopy-based leukemia-stroma coculture platform to systematically evaluate whether coculture models provide superior biological insights compared to monoculture studies. We developed a workflow suitable for high-throughput screening of primary leukemia and lymphoma samples cocultured with bone marrow stromal cells *ex vivo.* Our study elucidated that up to 50% of compounds, including BCR inhibitors, chemotherapeutics, and BET inhibitors, show reduced activity in the presence of bone marrow stromal cells. We observed very similar effects in lymphoid and myeloid malignancies, suggesting a disease-independent mechanism that mediates protection from drug-induced apoptosis. Carfilzomib and bortezomib for instance, two proteasome inhibitors, even lost their toxicity in CLL cells almost completely when cocultured with stromal cells. This observation might explain why proteasome inhibitors were clinically ineffective in CLL [48] patients. Our work complements current knowledge and underlines the importance to validate drug discoveries made in monoculture models in the context of different dimensions of the cancer microenvironment [19–21, 28–35].

Only few drugs were more active in coculture than in monoculture. JAK inhibitors reduced stroma-mediated protection in lymphoid and myeloid disease entities. The JAK-STAT pathway has been suggested as a key mediator of stromal protection [31, 32, 45, 46]. Our study confirmed the importance of the JAK-STAT signaling pathway for the crosstalk between leukemia and stromal cells. Although JAK inhibitors alone have low inhibitory activity, they could be used in combination with other clinically established drugs to reduce drug resistance in the bone marrow. Currently, JAK inhibitor combinations are being evaluated in clinical trials [49–51].

High-throughput *ex vivo* drug assays combined with molecular profiles of the tumor cells have demonstrated great potential to dissect genotype-phenotype relationships underlying drug response variation in cancer. Our data recapitulates clinically established genotype-drug response associations in CLL, suggesting that clinically relevant biology can be read out in short-term drug response assay *ex vivo*. Most of the identified drug-gene associations were consistent between mono- and coculture, but importantly the effect sizes of these associations in coculture were significantly reduced. Overall, our study demonstrates that monoculture drug assays represent an adequate discovery tool for drug-gene associations due to its lower complexity and higher sensitivity, whereas co-coculture systems are then necessary to estimate the *in vivo* relevance of a potential discovery in the context of the tumor microenvironment.

A limitation of our coculture model is the simplifying assumption that the mere presence of bone marrow-derived stromal cells is sufficient to reproduce the tumor microenvironment *ex vivo*. The bone marrow niche, however, represents a complex cellular system with many different cell types that interact with blood cancer cells and thus affect *in vivo* drug response. More complex coculture- and organoid systems [52–54] could address some of these limitations but our work suggests that simple coculture- or even monoculture models may produce informative drug response phenotypes.

## Acknowledgements

The authors thank all patients for the donation of cells. T.R. was supported by a fellowship of the German Federal Ministry of Education and Research (BMBF) and a physician scientist fellowship of the Medical Faculty of University Heidelberg. The authors gratefully acknowledge the data storage service SDS@hd supported by the Ministry of Science, Research, and the Arts Baden-Württemberg (MWK) and the German Research Foundation (DFG) through grant INST 35/1314-1 FUGG. T.R. was supported by a fellowship of the German Federal Ministry of Education and Research (BMBF) and a physician scientist fellowship of the Medical Faculty of University Heidelberg. S.D. was supported by the Else Kroener Physician Scientist Professorship, Heidelberg Research Centre for Molecular Medicine (HRCMM), an e:med BMBF junior group grant, and Deutsche Forschungsgemeinschaft (DFG). N.L. was supported by a Heidelberg School of Oncology (HSO2) fellowship from the National Center for Tumor Diseases (NCT) Heidelberg. The authors thank Benedikt Brors, Simon Anders, Beate Neumann, Sandrine Sander, Martina Seiffert, Christof von Kalle and Bernd Fischer for helpful discussions. The authors thank Angela Lenze, Tatjana Walter, Ximing Ding and Rainer Saffrich for technical support.

## Author contributions

S. A.H. conducted the screen. S.A.H., T.R., PM.B., C.K. and M.K. performed validation experiments. V.K. and S.A.H. performed computational analyses. E.C.S. performed data curation and J.L. processed N.G.S. data. N.L. collected patient data. C.L., P.D., C.M.T. and T. Z. provided guidance and feedback. S.A.H., V.K., T.R., and S.D. wrote the manuscript. S.A.H., V.K., and T.R. contributed equally to this work. S.D. and W.H. jointly supervised this work.

