## Supplementary Figures for "Comparing the value of mono- versus coculture for high-throughput compound screening in hematological malignancies"

### Supplementary Figure 1

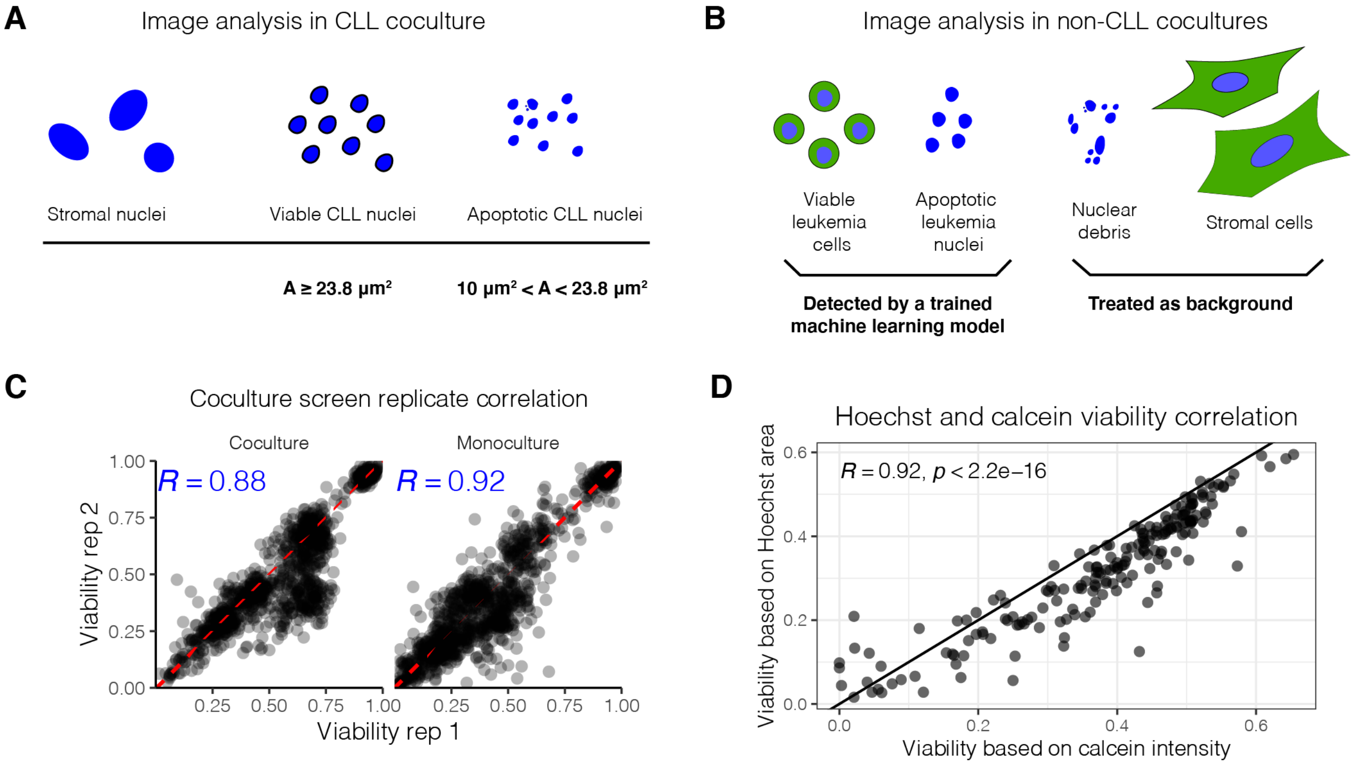


**Supplementary Figure 1. Reproducibility of the drug response readout.**

A) Image analysis in CLL coculture. B) Image analysis in non-CLL entities (AML, T-PLL, MCL, HCL). C) Replicate correlation in the coculture screen. D) Comparison of the viability readout based on CLL nucleus size and drug sensitivity readout with additional Calcein staining.

### Supplementary Figure 2

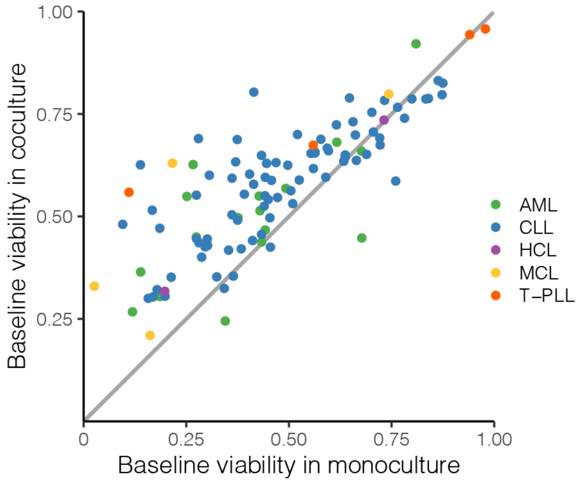


**Supplementary Figure 2. Spontaneous apoptosis in mono- and coculture.**

Leukemia cell viability without drug treatment. Each point corresponds to a leukemia sample (n = 108). X- and y- axes show median viabilities in mono- and coculture, with viability defined as the ratio of the viable cell count to the total cell count. The largest differences, and thus the strongest protection from spontaneous apoptosis, were observed in samples with low viability in monoculture.

### Supplementary Figure 3

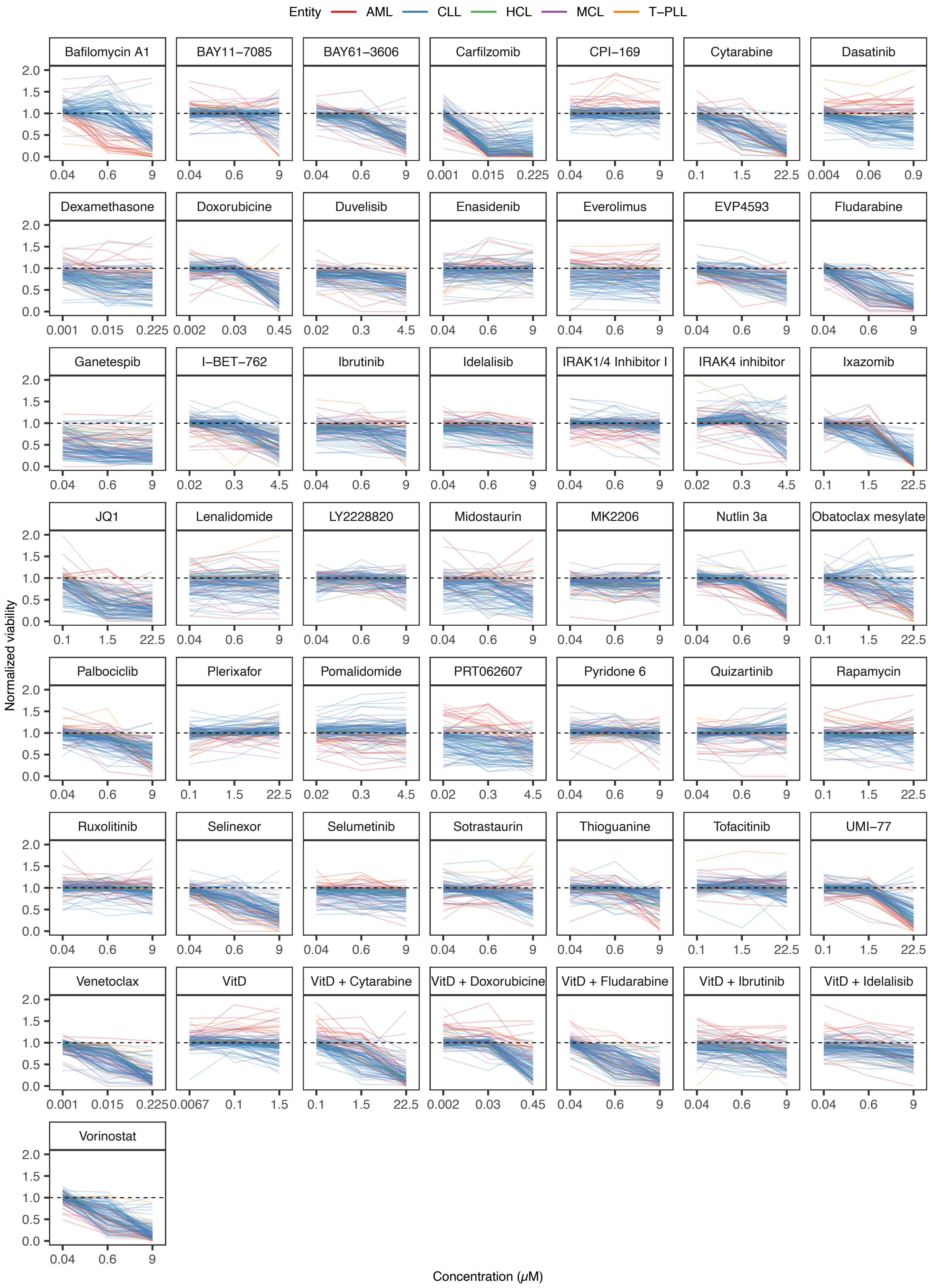


**Supplementary Figure 3. Drug response curves in monoculture.**

Line plots show normalized drug response curves in monoculture.

### Supplementary Figure 4

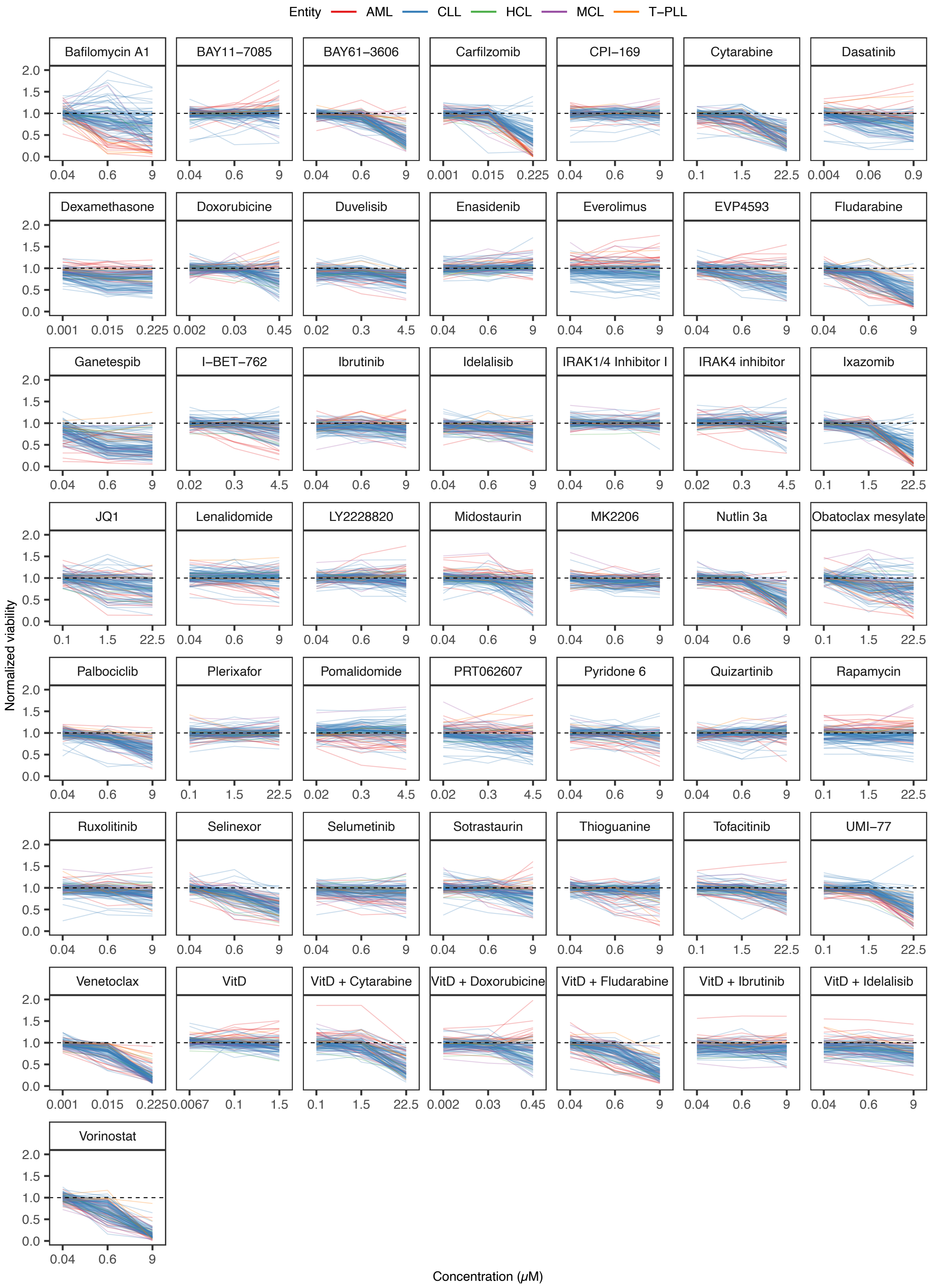


**Supplementary Figure 4. Drug response curves in coculture**

Line plots show normalized drug response curves in coculture.

### Supplementary Figure 5

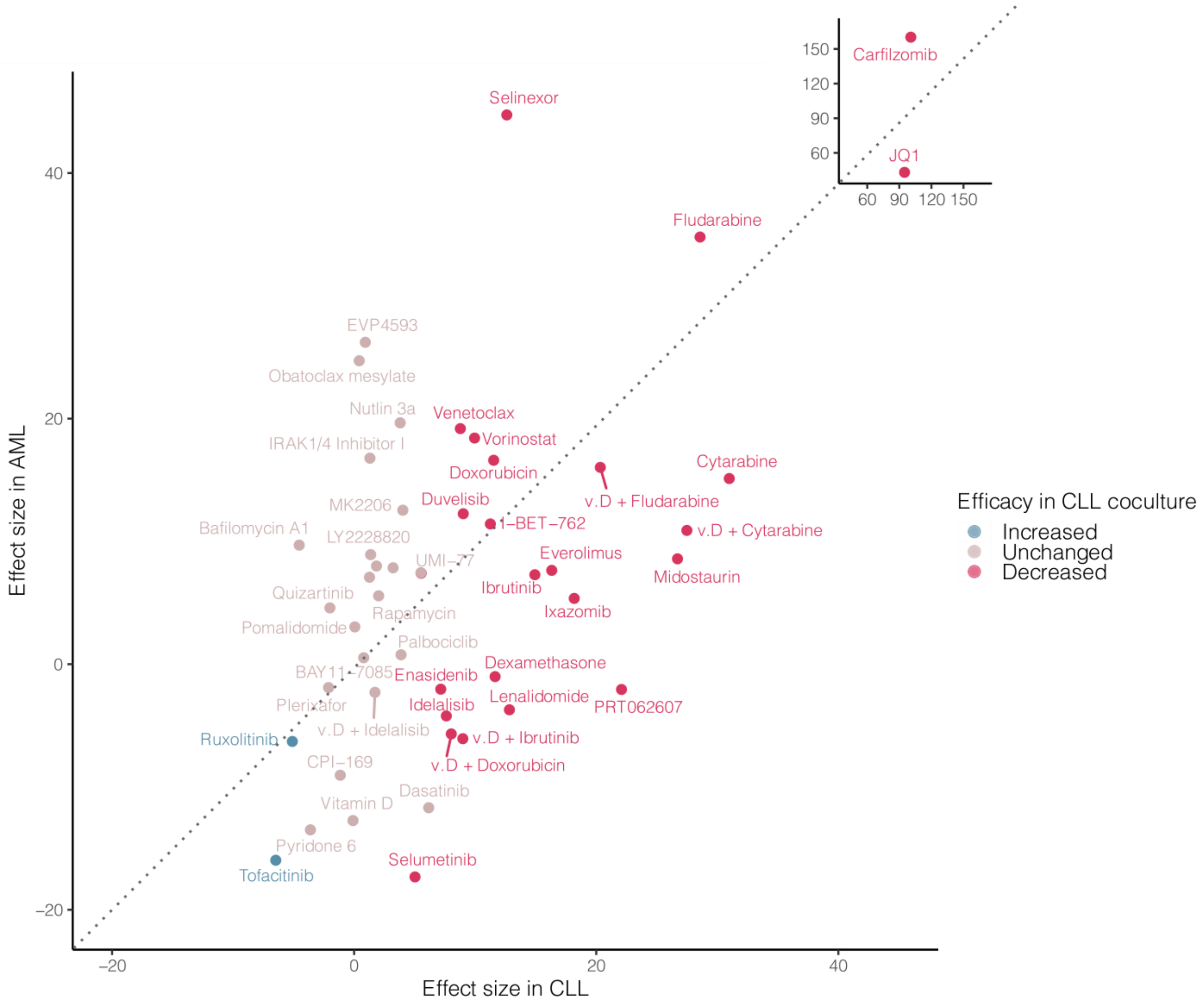


**Supplementary Figure 5. Effect size of efficacy changes in AML and CLL coculture.**
The scatter plot compares the effect sizes of drug efficacy changes in CLL-stroma coculture (x-axis) and AML-stroma coculture (y-axis). Points correspond to individual drugs probed in the screen. Point color indicates efficacy change in CLL-stroma coculture. (v. D. = vitamin D)

### Supplementary Figure 6

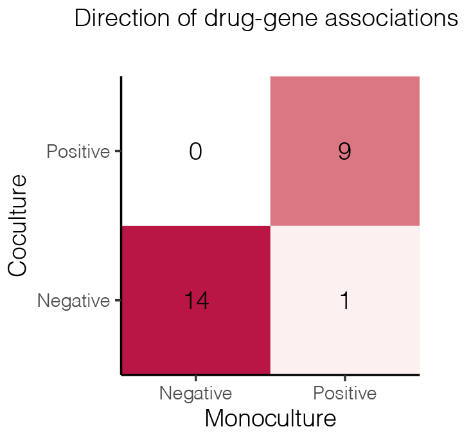


**Supplementary Figure 6. Gene associations in mono- and coculture.**

Contingency table comparing the direction of significant drug-gene associations (FDR < 0.1) in mono- and coculture.

### Supplementary Figure 7

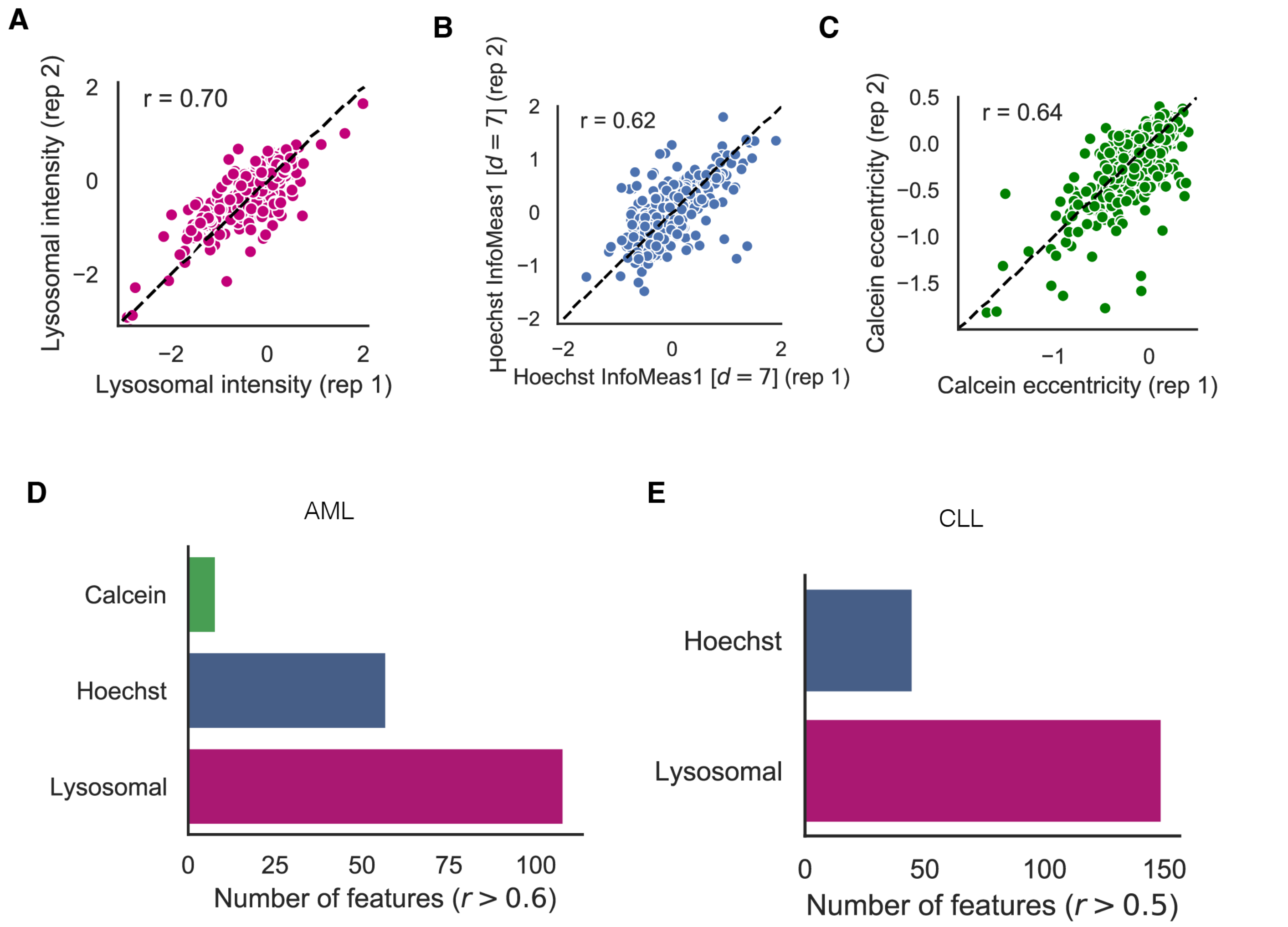


**Supplementary Figure 7. Morphological feature reproducibility.**

A)-C) Replicate correlations of lysosomal intensity, Hoechst InfoMeas1, and Calcein eccentricity. D) Number of reproducible image features (r > 0.6) in AML samples by color channel. E) Number of reproducible image features (r > 0.5) in CLL samples.

### Supplementary Figure 8

A

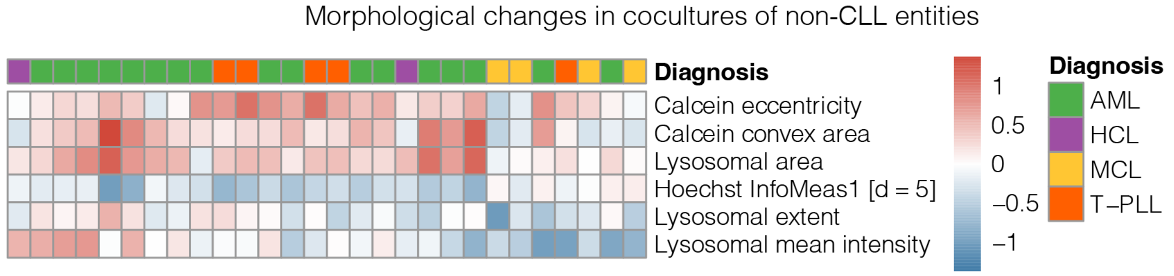


B

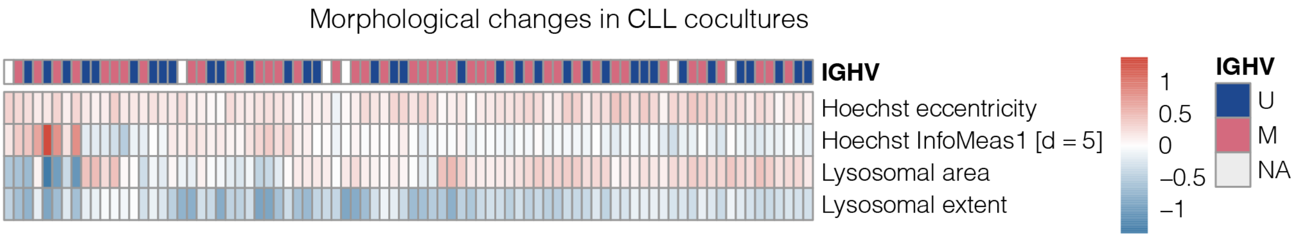


**Supplementary Figure 8. Morphological changes in coculture.**

A) Heatmap visualizing morphological changes in cocultures of non-CLL entities. Morphology changes were quantified as differences in medians of corresponding morphological features in coculture and monoculture. B) Morphological changes in cocultures of CLL samples annotated additionally with IGHV mutation status.

### Supplementary Figure 9

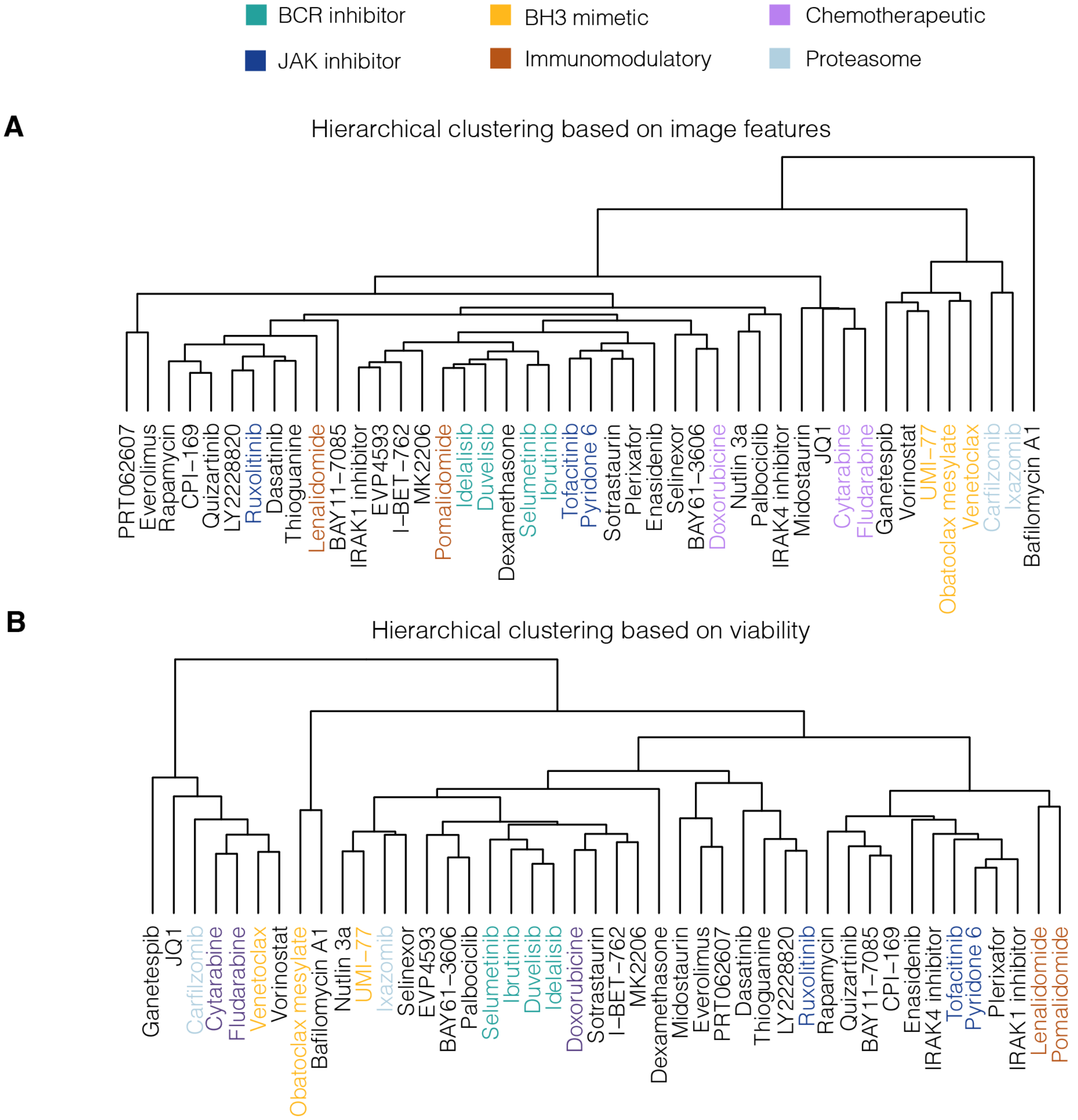


**Supplementary Figure 9. Hierarchical clustering of probed compounds.**
Hierarchical clustering based on A) image feature data alone and B) only viability. Text label color indicates drug class.
