## Supplementary Methods for "Comparing the value of mono- versus coculture for high-throughput compound screening in hematological malignancies"

*Compound screening of mono- and cocultures*

On the screening day, HS-5 stromal cells were detached and seeded at a density of $1x10^4$ cells/well in 20 µl into the columns with even numbers of CellCarrier-384 Ultra Microplates (Perkin Elmer). The high cell count enables even distribution of cells across the well. The columns with uneven numbers were filled with the same amount of medium. The cells were left at 37 °C for 3-4 hours to permit attachment. In the meantime, leukemia cells were thawed and allowed to recover, as described above. On each screening day either 5 or 10 patient samples were screened. Drug master plates were thawed and diluted with 96 µl serum free medium per well. 2.5 µl of drugs were transferred to the attached stroma cells, before adding 17.5 µL containing $2x10^4$ patient cells per well. The whole screen was carried out in DMEM (Thermo Fisher Scientific) supplemented with 10 % human serum (male AB, H6914-100ml Batch SLBT2873, Sigma-Aldrich), 1 % penicillin/streptomycin (Thermo Fisher Scientific) and 1 % glutamine (Thermo Fisher Scientific) at a final volume of 40 µL in the culture plates. Cells were incubated at 37 °C in a humidified atmosphere and 10 % $CO_2$ for 3 days.

*Processing of images (CLL)*

Images of CLL samples were processed using the image analysis software Harmony (Perkin Elmer). Stacks were processed by using maximum intensity projection. CLL nuclei were identified by segmentation of the Hoechst channel and separated from stroma cell nuclei based on the area of the nucleus. Results were exported and further analysis was conducted in the statistical programming language R (R Core Team, 2018). To assess whether a cell was alive or dead, the area of the nucleus of each individual cell was determined. When CLL cells die the nucleus condenses and, therefore, gets smaller and brighter. By plotting a histogram of nuclear area across all plates and patients we determined that a threshold of 23.8 $\mu m^2$ can distinguish the two populations of alive and dead cells most accurately. This method had previously been validated by concurrent staining with the viability dye Calcein AM (Supplementary Figure 1D). Using this threshold cells were classified into alive and dead, and the percentage of alive cells was calculated for each well.

*Image feature selection for compound profiling*

A non-redundant morphological feature set was constructed using a previously described method [2]. Briefly, an initial set of 2-3 features was provided. At each iteration, all other features were regressed against the set of selected features and the feature with the highest replicate correlation between regression residuals was added. The iterative procedure continued until the number of features with positive correlations between residuals was exceeding the number of those with negative correlations. The selected features used for compound profiling are provided in Supplementary Tables 6-7.

*Quantification and visualization of morphological changes*

Median values were estimated for each morphological feature in every probed condition. The difference of medians ($m_C$ - $m_M$) in coculture and monoculture was computed to quantify stroma-induced morphological changes in each sample. For mathematical definition of morphological properties see Supplementary Table 8. Coculture and monoculture cell populations of each sample were jointly embedded using t-SNE [1]. The t-SNE algorithm was applied to the first 20 principal components (PCs) of single-cell morphological feature data.

*Compound profiling and hierarchical clustering*

Selected morphological features were used to generate compound profiles by aggregating the observed features across all screened samples in both culture conditions. These image-based profiles were used for hierarchical clustering of probed compounds. Drug-drug correlations of image-based profiles were used to measure drug similarities independently in mono- and coculture.

*IGHV status analysis*

For the analysis of IGHV status RNA was isolated from $1x10^7$ PBMCs and cDNA was synthesized via reverse transcription. Subsequent PCR reactions and analyses were based on the protocol published by Szanaski and Bahler [3] with minor modifications. The AmpliTaq Gold DNA polymerase (Thermo Fisher Scientific) was used for PCR reactions. Amplification of VH1-, VH3- and VH4- segments was done in single reactions whereas amplification for VH2, VH3-21, VH5 and VH6-segments was done in a multiplex fashion as previously described (Primers for the individual PCRs: PCR1: VH1, JH, JH-1; PCR2: VH3, JH, JH-1; PCR3: VH4, JH, JH-1; PCR4: VH2, VH3-21, VH5, VH6, JH, JH-1) [3]. The following PCR program was used: initial denaturation (94 °C, 2 min), 40 cycles of denaturation, annealing and elongation (94 °C, 20 sec; 52 °C, 10 sec; 72 °C, 30 sec) and final elongation (72 °C, 2 min). Sanger Sequencing (GATC Biotech) was performed on the PCR products using the appropriate forward and the JH-1 reverse primer. In the multiplex PCR reaction both JH-rev and JH-1 rev were used for sequencing. Forward and reverse sequencing results were aligned. The IMGT/V-Quest-Database was used for finding the closest matching germline VH-sequence and identifying the mutation status, i.e., the percentage of sequence identity, of the VH-segment determined.

*Panel sequencing of CLL samples*

Analysis of gene mutations of the genes NOTCH1, SF3B1, ATM, TP53, RPS15, BIRC3, MYD88, FBXW7, POT1, XPO1, NFKBIE, EGR2 and BRAF was performed. A customized Illumina™ TruSeq Custom Amplicon (TSCA) panel with two independent primer sets for a redundant coverage of genes was designed. The selection of targets comprised the 11 most frequently mutated genes in CLL identified via unbiased whole exome sequencing of 528 CLL patients [4]. For the following genes the full gene was covered: ATM, BIRC3, EGR2, FBXW7, MYD88, NFKBIE, POT1 and TP53. For the following genes the most affected exons were covered: BRAF (exons 11-18), NOTCH1 (exon 34 +3'UTR), RPS15 (exons 3-4), SF3B1 (exons 14-16) and XPO1 (exons 14-17). Library preparation was performed using TruSeq Custom Amplicon Assay Kit v1.5 including extension and ligation steps between custom probes. Samples were indexed, pooled, and loaded on an Illumina MiSeq flow cell in 32 sample batches.

Analysis was performed using BWA, Samtools (alignment), and Varscan (variant calling and annotation) [5, 6]. Current databases (COSMIC, 1000G, dbSNP, ClinVar) [7-10] were considered to evaluate variants above a threshold of 5 % mean variant allele fraction (VAF) as pathogenic/nonpathogenic. Only mutations occurring at an allele frequency of more than 20 % and in at least three patients were considered.

*Validation of screening results in primary MSC cocultures*

Primary mesenchymal stromal cells (MSCs) derived from three different healthy donors were used. 1000 cells/well were seeded in 96-well glass bottom plates (zell-kontakt GmbH) in

MSCGM$^{TM}$ Mesenchymal Stem Cell Growth Bulletkit Medium (Lonza). The plates were cultured in a humidified atmosphere at 37 °C and 5 % $CO_2$ for 2 days to allow MSCs to adhere and recover. CLL cells were thawed, allowed to recover in medium for 3 hours and filtered through a 40 µm cell strainer (Sarstedt) to get rid of dead cells. The medium was removed from MSCs and $2x10^5$ CLL cells/well added to the stroma cells in Bulletkit medium (Lonza). Apart from cocultures, monocultures of CLL cells were established. Cultures were treated with 1.5 µM JQ1, 0.6 µM Fludarabine, 22.5 µM tofacitinib, 9 µM ruxolitinib or solvent control (DMSO; SERVA Electrophoresis GmbH) and incubated in a humidified atmosphere at 37 °C and 5 % $CO_2$ for 3 days. Each condition was assessed in technical duplicates. The cultures were stained with Hoechst 33342 (4 µg/ml, Invitrogen), Calcein AM (1 µM, Invitrogen), PI (5 µg/ml, Sigma-Aldrich) and lysosomal dye NIR (1 µl/ml, Abcam) and the whole wells were imaged on an Opera Phenix microscope (Perkin Elmer) with a 10x objective in confocal mode. Using the image analysis software Harmony (Perkin Elmer), CLL cells were segmented based on Hoechst signal and Calcein and PI intensities were measured. All further analysis steps were conducted in R (R Core Team, 2018). Cells with a Calcein intensity above a certain threshold and with PI intensities below a certain threshold were classified as alive. To avoid a possible influence of phagocytosis on relative percentages of alive cells [11], absolute counts of alive CLL cells were used for all further analyses. These viabilities were normalized by division through viabilities in DMSO controls.

*Western blot analysis*

Samples for Western Blot were prepared by washing once with ice-cold PBS (Thermo Fisher Scientific) and lysed in 100 µl RIPA buffer (Sigma-Aldrich) containing PhosSTOP (Sigma-Aldrich) and cOmplete, Mini Protease Inhibitor Cocktail (Sigma-Aldrich). After incubation on ice for 30 min the samples were centrifuged at 15,000 g for 20 min at 4 °C. The supernatant was aliquoted and frozen at -80 °C until use. Samples were run on 10 % acrylamide gels (SERVA Electrophoresis GmbH) at 45 mA. As marker, the dual color Precision Plus (Bio-Rad Laboratories) ladder was used. Transfer to PVDF membranes (Thermo Fisher Scientific) was performed at 400 mA. Primary antibodies, as listed in the main part of the manuscript, were incubated overnight.
